# Age and Generation-Based Model of Metastatic Cancer: From Micrometastases to Macrometastases

**DOI:** 10.1101/2025.06.26.661673

**Authors:** Panagiotis Gavriliadis, Georgios Lolas, Themis Matsoukas

## Abstract

Metastasis remains the decisive event in most solid cancers, yet the mathematical tools we use to describe it are still largely tied to tumor size. In the curent work, we recast metastatic spread in terms of tumor age and generational ancestry, turning an implicit notion into the main organizing principle of the model. Building on the classical Iwata–Kawasaki–Shigesada (IKS) framework, we construct a transformation from size to age and derive a hierarchy of integral equations indexed by metastatic generation. Our reformulation yields closed-form expressions for first-generation metastases and simple one-dimensional recursive integrals for higher generations, avoiding direct numerical solution of the original size-structured partial differential equation (PDE) with its nonlocal boundary condition. Using Gompertzian growth and power-law metastatic emission, we show that the age–generation model reproduces IKS predictions over clinically relevant time scales while offering improved numerical stability and interpretability.

The generational decomposition reveals a robust pattern: lesions seeded directly from the primary dominate early in the disease course, whereas successively younger generations, emitted by existing metastases, come to dominate the total lesion count at small sizes, leaving older generations to occupy the macroscopic tail of the distribution. Introducing an explicit detection threshold naturally separates a small number of radiologically visible macrometastases from a much larger, unseen pool of micrometastases. Together, these results provide a transparent and computationally efficient framework that links primary-tumor growth, metastatic seeding across generations, and the hidden microscopic burden that underlies clinical presentation.

## 1. Introduction

Cancer is not a single disease but a heterogeneous collection of malignancies that can arise in virtually any tissue, follow distinct evolutionary paths, and respond very differently to therapy [1]. Despite tumor diversity, most cancerrelated deaths share a common proximal cause: metastatic dissemination [2]. Solid tumors such as breast, lung, pancreas, prostate, colorectal cancer and melanoma may remain clinically silent or locally confined for long periods, yet once malignant cells successfully seed distant organs and establish secondary lesions, prognosis deteriorates sharply [3, 4].

In both preclinical and clinical settings, tumors are monitoblue almost exclusively through their size. Radiological imaging provides diameters or volumes of detectable lesions, while in animal models calipers and bioluminescence or fluorescence signals play a similar role. Tumor size has thus become the de facto proxy for tumor burden and metastatic progression. Cancer size-based description is convenient but incomplete. It conflates two fundamentally different aspects of tumor dynamics: intrinsic growth kinetics and the time elapsed since a lesion was initiated. A large tumor may be old and slowly proliferating or young and aggressively expanding; both scenarios may carry very different risks and therapeutic implications, yet they can be indistinguishable when assessed only through size.

The latent quantity that resolves this ambiguity is tumor age, understood as the time elapsed since a lesion was initiated at some reference size (for example a single cell or a minimal detectable cluster of cancer cells). Under standard growth laws used in oncology (exponential, power-law, logistic, Gompertz) and within the regime of untreated or minimally treated disease before strong necrosis or therapy-induced shrinkage, tumor size typically evolves monotonically. Along each growth trajectory, size and age move in lockstep: to each biologically admissible size there corresponds a unique age and vice versa. Existing mathematical descriptions of metastatic spread, however, have kept this size–age link implicit and have almost exclusively adopted a size-based perspective (see reviews from [5, 6, 7, 8, 9, 10]).

In the current work we turn tumor age from an implicit variable into the central organising coordinate of the metastatic process. Building on the classical size-based model of Iwata, Kawasaki and Shigesada (IKS) [11], we construct an explicit transformation between size and age and use it to reparametrise metastatic dynamics in terms of tumor age and metastatic generation. First-generation metastases are those seeded directly by the primary tumor, second-generation metastases are seeded by the first generation, and so on. Such an approach leads to a recursive age and generation-based representation of metastatic dissemination that bypasses the size based IKS formulation.

### 1.1. A sneak peek on the IKS model

A landmark advancement in mathematical modeling of metastasis was introduced by Iwata, Kawasaki, and Shigesada [11]. The IKS model is based on a variant of the McKendrick-von Foerster equation [12, 13], a structured population model commonly utilized in epidemiology and ecology, to track the dynamics of age-structured populations [14]. In cancer biology, the IKS model established a mathematical connection between tumor growth and metastatic burden, a relationship that is often difficult to infer from clinical observations. In the IKS model, tumors of size *x* grow at a rate of *g*(*x*) and metastasize at a rate of *β*(*x*), with each metastasis producing new tumors of size *x* = 1, which then grow at a rate of *g*(*x*) and emit metastases at a rate of *β*(*x*) (see Fig. 1). These rates are given by [11]

**Figure 1.**
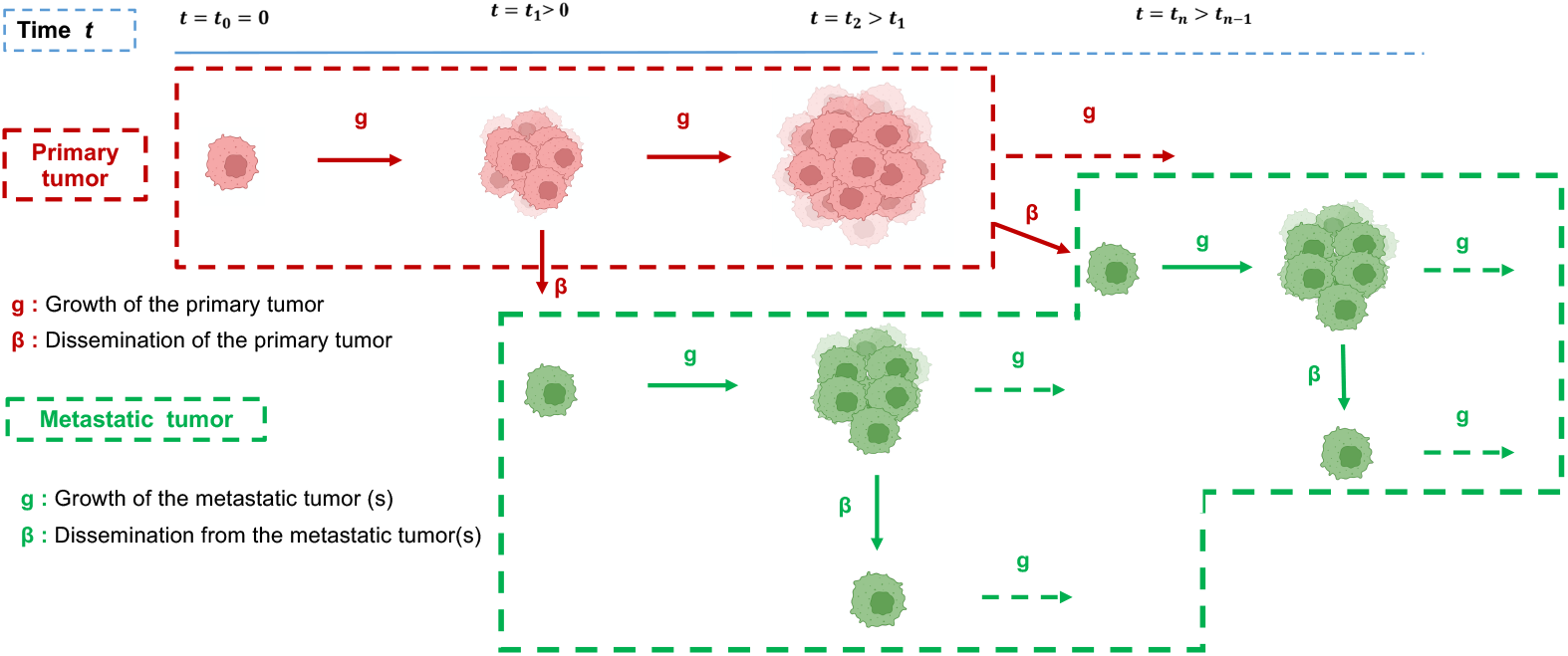
A cartoon depiction of the IKS model illustrating primary tumor growth and metastatic dissemination. The primary tumor emits metastatic cells that subsequently form secondary metastatic tumors. Growth (*g*) and dissemination rates (*β*) for both primary and metastatic tumors are indicated.

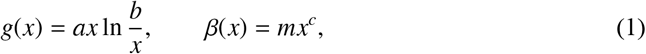

withe *a, b, m* and *c* constants. The above expression for *g*(*x*) is the so-called Gompertz function. In general, *g* : [*x*_0_ = 0, *x*_1_ = *b*] → [0, +∞) is continuous, *g >* 0 for *x > x*_0_, *g*(*x*_1_) = 0 and 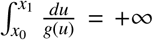. Regarding the birth function, we assume that *β* ∈ *L*^∞^([*x*_0_, *x*_1_]) and *β*(*x*) *>* 0 on [*x*_0_, *x*_1_]. The distribution density of the metastatic tumors, *ρ*(*x*; *t*), is governed by the transport equation [11]

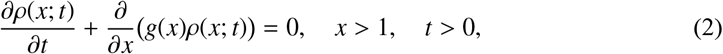

along with the boundary condition

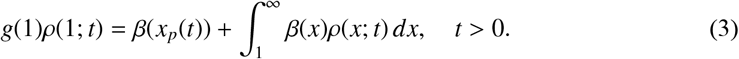

which describes the number of “elementary tumors” (metastatic embryos of size *x*_0_ = 1) generated from the primary tumor, whose size at time *t* is *x*_*p*_(*t*), and from all metastatic tumors. Equation (2) is supplemented with the initial condition

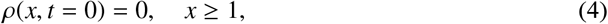

which states that no metastatic tumor exists at the initial time (see Appendix A for a formal derivation).

The IKS model describes metastatic growth via a size-structured PDE for which explicit analytic formulas are available only in the asymptotic regime *t* → ∞ [11, 15], so finite-time solutions must be obtained numerically despite the challenges posed by a nonlocal boundary condition and biologically relevant tumor sizes spanning from a single cell to more than 10^10^ cells over decades [16, 17, 18]. Tumor size difficulties have been tackled by logarithmic domain compression and high-order Runge–Kutta schemes [16], by convergent characteristic-based methods inspired by general size-structured transport solvers [15, 19], and by reformulating the model as a system of Volterra integral equations [20], a framework on which Bulai and co-authors [21, 22] later developed specialized solvers in one and two dimensions.

### 1.2. Extensions of the IKS model: biological and clinical developments

Beyond numerical optimization, the IKS model has served as a foundation for biological and clinical extensions: Benzekry and co-workers [23, 24, 25] linked size-structured metastatic dynamics with angiogenesis through an ODE-based extension of the Hahnfeldt model and later related presurgical tumor volume to postsurgical metastatic burden, while Barbolosi *et al*. [26] integrated drug efficacy into metastatic burden overpredict, Bilous *et al*. [27] incorporated patient-specific data, treatment effects and dormancy phases to describe brain metastases in NSCLC, and Liu and Wang [18] proposed a unified size-structured PDE model enabling optimal bang–bang chemotherapy scheduling. Further refinements include mathematical analyses of the IKS model and its nonlocal boundary condition [28], a Monte Carlo formulation of the IKS framework [29], biologically informed models with dormancy, secondary emissions and interindividual variability validated in mice [30], clinically calibrated extensions including systemic therapy for lung cancer patients [31, 32], zebrafish-based population-balance models [33], IKS-based multiscale parametric models linked to quantitative imaging biomarkers in multiple myeloma [34] and the integration of IKS-type mechanisms into machine-learning frameworks for perioperative and neoadjuvant treatment evaluation [35].

### 1.3. A generational perspective

As Devys and colleagues noted [16], when the population balance is formulated from a generational perspective–offspring of the primary tumor, offspring-of-the-offspring of the primary tumor, and so on–the contribution of the tumor distribution in each generation to the total is additive. The distribution of each generation is calculated by a transport equation similar to Eq. (2) with Eq. (3) replaced by a recursion. Here, we take the idea of Devys *et al* [16] one step further by reformulating the metastatic process in terms of tumor age rather than size. We then demonstrate that the effects of each generation can be distinctly separated and can be calculated through recursive one-dimensional integrals that eliminate the need to solve any partial differential equations. The remainder of the paper is organized as follows: In Section 2 we formulate the distribution in terms of the tumor age. In Section 3 we derive the distribution of tumors produced solely by emissions from the primary tumor (first generation of emissions), and then we extend the calculation to all subsequent generations, resulting in a recursion that involves only integrals over the distribution of the previous generation. In Section 4 we discuss the extension to growth and emission rates that vary among generations while in Section 5 we apply our methodology to the IKS model and compare the results to the standard solution of the transport equation. We present our findings in Section 6 and discuss them in Section 7. Finally, we provide our concluding remarks in Section 8.

## 2. Age-based framework for metastatic tumours

### 2.1. Age–size relationship

The age *τ* of a tumor is the interval of time for a tumor of initial size *x* = 1 to reach size *x* = *x*(*τ*). Accordingly, the relationship between size and age is simply given by the growth

equation:

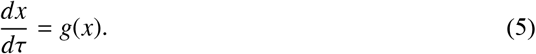

As long as the growth rate is positive the relationship between *x* and *τ* is monotonic and either variable can be expressed in terms of the other,

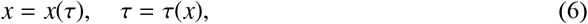

with *x*(0) = 1, *τ*(1) = 0. This relationship allows us to describe the state of the tumor in terms of either size or age in completely equivalent terms.

The size–to–age transformation is valid precisely when the net growth rate *g*(*x*) remains positive, so that tumor trajectories are strictly increasing and the mapping between size and age is one-to-one. Biologically, this corresponds to the regime of untreated or minimally treated solid tumors prior to pronounced necrosis or therapy-induced shrinkage, and is not intended to cover late-stage disease with cycles of regression and regrowth where a more general, nonmonotone age–size relation would be required.

### 2.2. Age distribution

The distribution of tumors in terms of size *x* is *ρ*(*x*; *t*), such that the number of tumors *dN*(*t*) at time *t*, whose size is between *x* and *x* + *dx*, is:

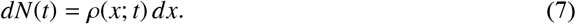

Let *φ*(*τ*; *t*) be the distribution of tumors in terms of their age *τ*. In terms of this distribution, the number *dN*(*t*) of tumors at time *t*, whose age is between *τ* and *τ* + *dτ* is:

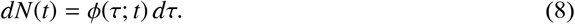

Combining these results and using *g* = *dx/dτ* from Eq. (5) we obtain the relationship between *ρ*(*x*; *t*) and *φ*(*τ*; *t*):

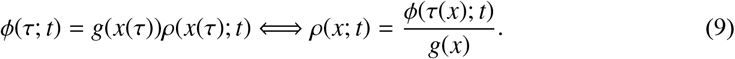

Just as *x* and *τ* are equivalent ways to describe the state of a tumor, *ρ*(*x*; *t*) and *φ*(*τ*; *t*) are equivalent descriptions of the distribution of tumors. We note that *φ*(*τ*; *t*) receives nonzero values only in the interval *τ* = 0 to *τ* = *t*, since the maximum age at time *t* is *t*, corresponding to the age of the primary tumor. Similarly, the distribution *ρ*(*x*; *t*) receives nonzero values in the interval *x* = 1 to *x*_*p*_(*t*), where *x*_*p*_(*t*) is the size of the primary tumor at time *t*.

### 2.3. Production of tumors

All metastatic tumors at time *t* with age *τ* originate from the total emissions at time *t* − *τ*. Accordingly, the number of these tumors is equal to the number of emissions from all tumors at time *t* − *τ*, primary and metastatic. The emission rate from the primary tumor is *Q*(*t* − *τ*), where *Q*(*t*) is given by

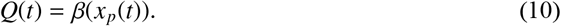

The emission rate from metastatic tumors of age 0 ≤ *τ*’ ≤ *t* − *τ* is *φ*(*τ*’; *t*)*β*(*x*(*τ*’)), and their total emission rate is the integral of this quantity over all *τ*’. The metastatic distribution *φ*(*τ*; *t*) is the sum of the two contributions:

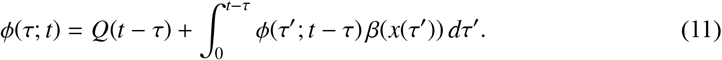

Applying Eq. (9) to the above result we obtain an analogous equation for the distribution of tumor sizes:

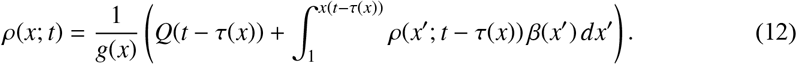

With Eqs. (11) and (12) we have obtained the distribution of tumors in age and size, respectively, at any time *t* in terms of the corresponding distribution at all previous times. Next we will obtain closed form solutions for these distributions.

## 3. Hierarchical decomposition of emissions

### 3.1. First-order emissions

In the early phase of cancer evolution, when individual lesions are still small and shed metastatic cells only very slowly, the metastatic burden is dominated by the primary tumor, which is both the largest lesion and remains so throughout this period. In the first-order approximation we ignore all other metastases except for those from the primary tumor; dropping the integral in Eq. (11) (since for *k* = 1 there are no parent metastases).

Indicating the contribution of the first-order approximation by *φ*_1_, the result is

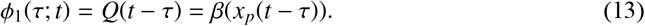

Using Eq. (9) we obtain the corresponding size distribution of primary emissions:

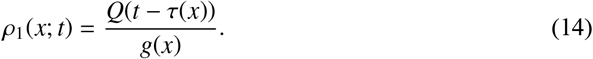

### 3.2. Higher order emissions: The generational viewpoint

Metastatic emissions obey a hierarchy that allows the decomposition of their effects in a systematic manner. This hierarchy is illustrated in Fig. 2 as a branching graph in discrete time steps. The main stem of the graph (Fig. 2a) represents the primary tumor. In the absence of emissions the tumor grows in size in each time step. Physical time is shown in blue and the age of the tumor is shown in grey. Primary metastases (metastases originating from the primary tumor) constitute generation *k* = 1 and are shown in Fig. 2b with their corresponding age in magenta. Secondary emissions originate from secondary tumors and constitute generation *k* = 2, shown in blue in Fig. 2c.

**Figure 2.**
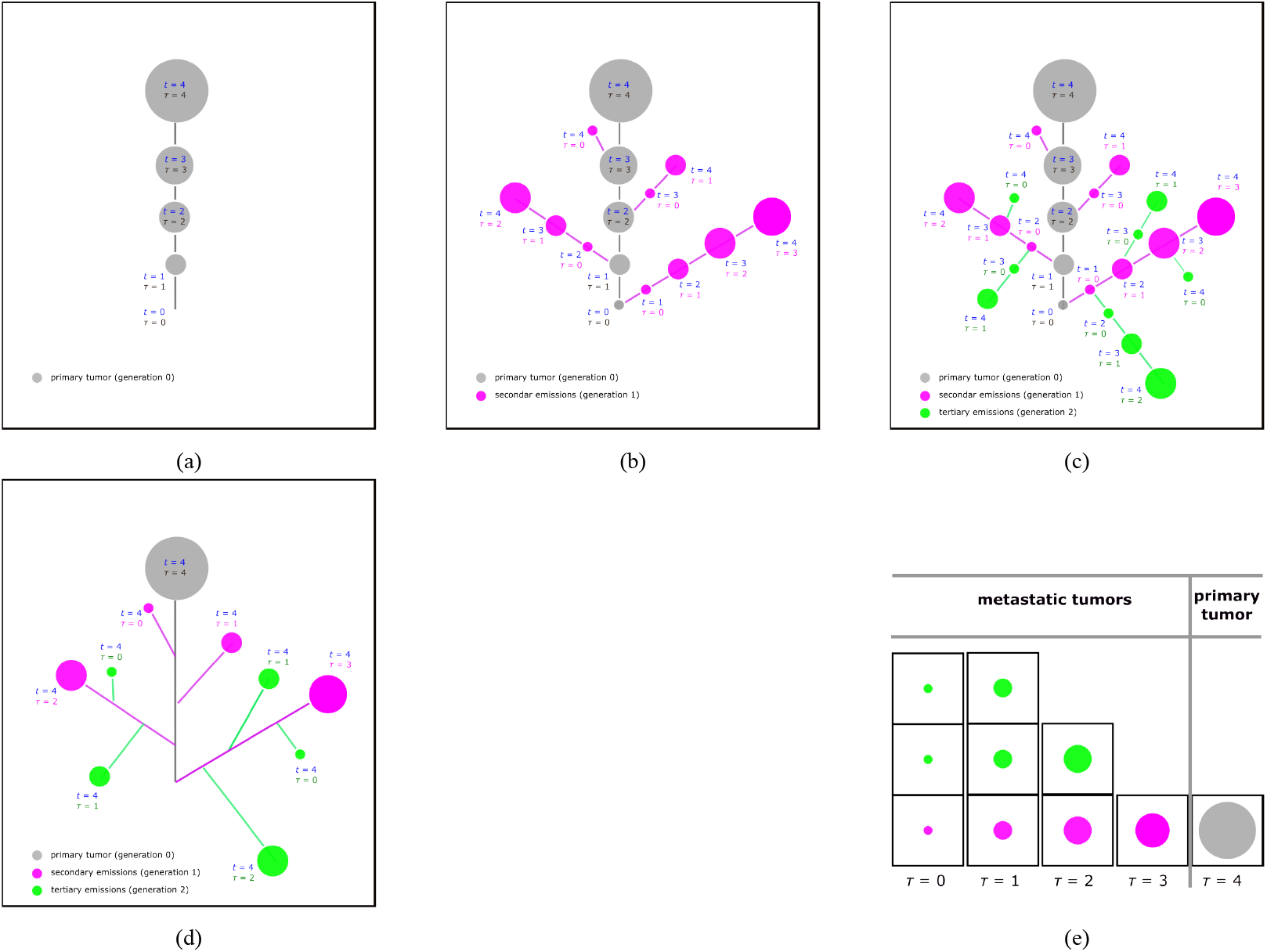
Hierarchy of Tumors. (a) Evolution of primary tumor (generation 0) in discrete time steps δ*t* = 1 up to *t* = 4δ*t*. Physical time is indicated in blue, the age of the tumor is indicated in gray (b) Evolution of secondary emissions (generation 1) up to *t* = 4δ*t*. The age of the tumor is indicated in magenta. (c) Evolution of tertiary emissions (generation 2) up to *t* = 4δ*t*. The age of the tumor is shown in blue. (d) Population of tumors at *t* = 4δ*t*. (e) Distribution of tumors at *t* = 4δ*t*. There are three metastatic tumors of age τ = 0, three of age τ = δ*t*, two of age τ = 2δ*t* and one of age τ = 3δ*t*. The primary tumor at *t* = 4δ*t* is of age τ = 4δ*t*. Here the discrete steps δ*t* are used only for illustration; in the model itself, physical time *t* is continuous.

Extension to higher order emissions is straightforward. Each new generation *k* branches out from branches of its parent generation *k* − 1. Physical time is equal to the number of steps to reach a particular tumor while the age of the tumor is the number of steps starting from the beginning of the branch to which the tumor is attached. Every branch is a clone of the primary tumor and it is transposed in physical time by the birthtime of the branch. The state of the tumor population at time *t* is the ensemble of all tumors that correspond to the same physical time *t*.

This is shown schematically in Fig. 2d, which illustrates the population at discrete time *t* = 4. The age distribution of the population is illustrated in Fig. 2e. It is clear from this illustration that the age distribution is the sum of the age distribution within each generation. That is,

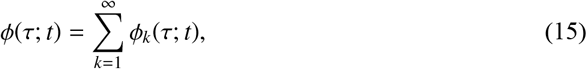

where φ_*k*_(τ; *t*) is the age distribution of generation *k* of metastatic emissions at time *t*. Since φ and ρ are proportional to each other, the size distribution is given by a similar series:

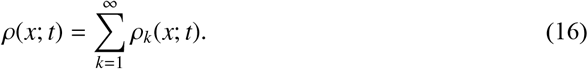

At fixed time *t* the contribution of each subsequent generation to the distribution of tumors decreases, as older and thus bigger tumors emit at a substantially higher rate. We may then construct the *m*-order approximation by retaining only *m* terms in the series:

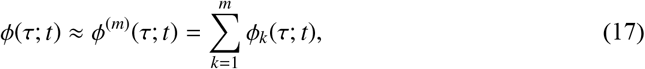

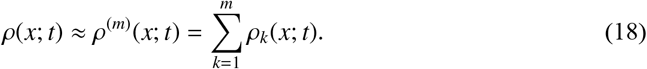

Next we obtain equations for the distribution in each generation.

### 3.3. Recursive solution

All metastatic tumors in generation *k* whose age at time *t* is *τ* grew from newly born metastatic seeds were born at time *t* − *τ* from all metastases of order *k* − 1. Integrating the emission rate from parents of all possible ages, noting that only parents with age *τ*’ ≤ *t* − *τ* produce branches whose age at *t* − *τ* is *τ*, we have

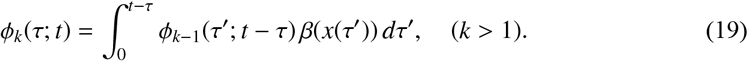

This forms an equation for the distribution of metastatic cancers of generation *k* in terms of the distribution of the parent generation *k* − 1. Along with Eq. (13) for *k* = 1 we have a closed system that can be solved recursively for any generation.

To obtain an analogous equation for the size distribution of tumors we first note that by direct analogy to Eq. (9), the relationship between *ρ*_*k*_ and *φ*_*k*_ is

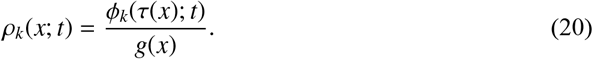

We apply the above along with *φ*_*k*−1_(*τ*’; *t* − *τ*)*dτ*’ = *ρ*_*k*−1_(*x*’; *t* − *τ*)*dx*’ to Eq. (19) to obtain the size distribution in generation *k*:

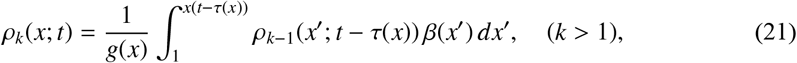

where *x*(*t* − *τ*(*x*)) is the size of the (*k* − 1)order metastatic tumor at time *t* − *τ*(*x*). Along with Eq. (14), we have a closed system that can be solved recursively for the size distribution in any generation. Our generational solution is presented in Fig. 3.

**Figure 3.**
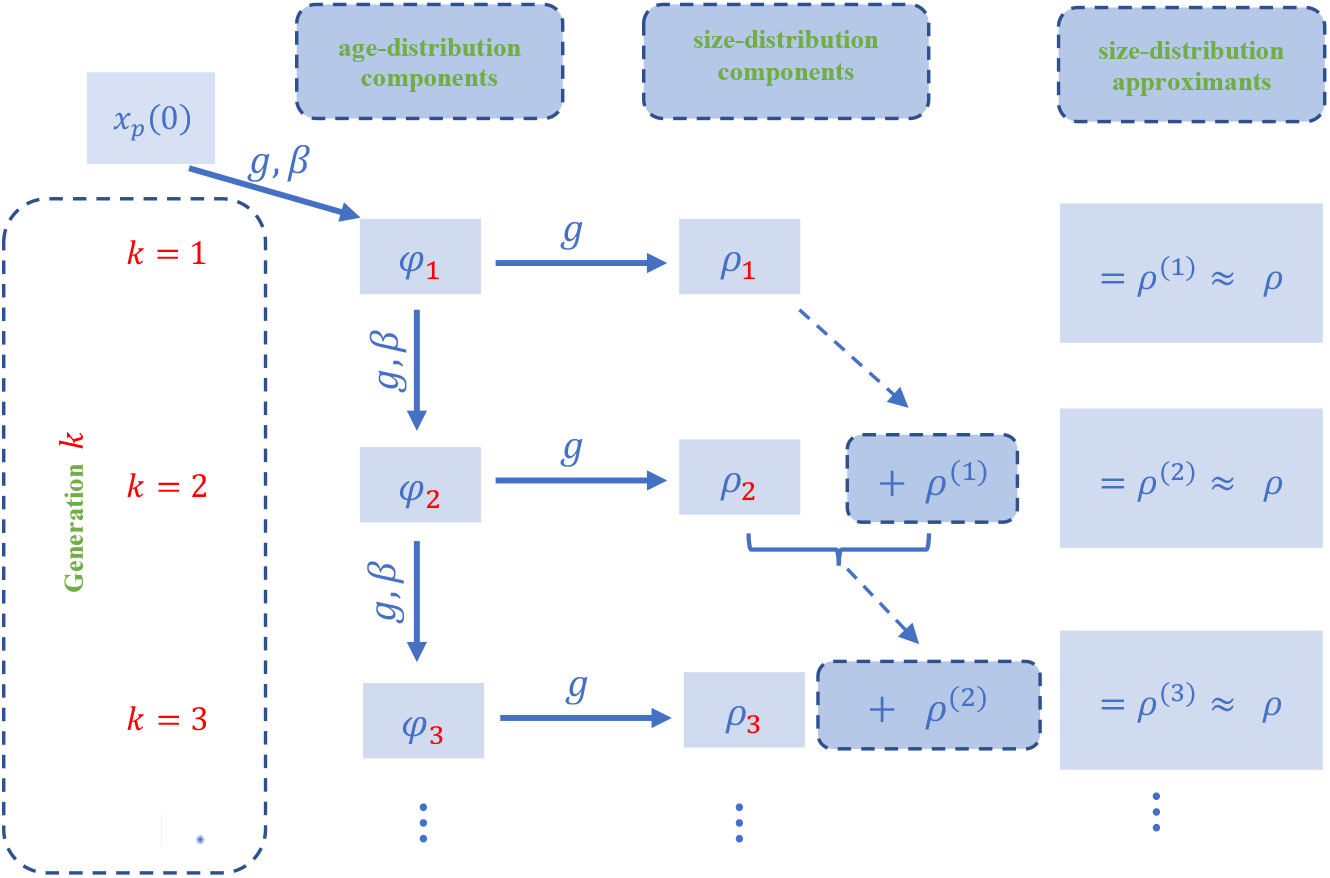
Generation–resolved construction of the metastatic size distribution. For each generation *k* = 1, 2, 3, …, the age–distribution *φ*_*k*_(*τ*; *t*) is first obtained from the recursive relations in age space. The arrows indicate how each *φ*_*k*_ is mapped, via the age–size transformation and the growth law, to the corresponding size–distribution component *ρ*_*k*_(*x*; *t*). By summing these components over generations one obtains successive *k*th–order approximations 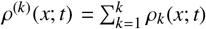 of the full metastatic size distribution *ρ*(*x*; *t*). The diagram is schematic and is meant to highlight the recursive passage from generation–resolved age densities to generation–resolved size densities and, finally, to their cumulative approximation of the total size distribution.

## 4. Extension to generation-dependent rates

Previously, we assumed that all generations shared the same growth law *g*(*x*) and emission rate *β*(*x*). We now allow each generation *k* to have its own functions *g*_*k*_(*x*) and *β*_*k*_(*x*), so that different generations can grow and emit metastases at different rates. This generalization makes it possible to describe biological heterogeneity, environmental differences between primary and secondary sites, or evolutionary changes in tumor aggressiveness across generations.

For each generation *k*, let *x*_*k*_ = *x*_*k*_(*τ*) denote the size of a tumor of generation *k* as a function of its age *τ*. It satisfies the growth law

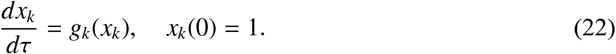

The inverse function *τ*_*k*_ = *τ*_*k*_(*x*) gives the age corresponding to a tumor of generation *k* that has reached size *x* under its specific growth law; in other words, *τ*_*k*_(*x*) is the time needed for a generation-*k* tumor to grow from the reference size 1 to the size *x*. As in the base model, each growth function *g*_*k*_(*x*) is assumed to remain positive and monotonic in *x*, ensuring that the mapping between size and age is one-to-one.

By direct analogy to Eq. (19) the age distribution of generation *k* (*k >* 1) at time *t*, is

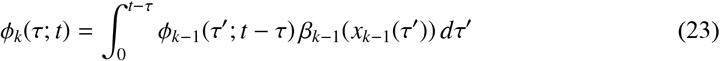

and, similarly, by analogy to Eq. (21) the size distribution in generation *k* (*k >* 1), is

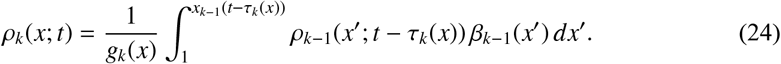

The straightforward conversion of Eqs. (19) and (21) into (23) and (24), respectively, is possible because a fundamental property of the metastatic model is that each new generation is completely determined by its parent generation–there is no interference from any other branches. For *k* = 1, Eqs. (13 14) remain valid.

## 5. Numerical implementation with Gompertzian growth and power-law metastatic emission

We now apply the recursive framework to Gompertz growth with a power-law emission, as in the original IKS model. In this setting, the first generation admits a closed-form solution and higher generations are computed from simple one-dimensional integrals, while all IKS parameters and biological scalings are preserved. The growth and metastatic emission rates are given by

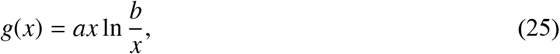

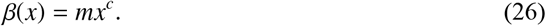

For the purposes of numerical calculations we will use the following parameters,

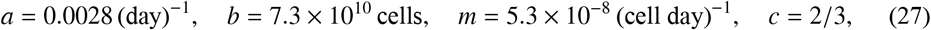

which were used by Iwata *et al* [11] based on clinical observations of metastatic hepatocarcinoma.

### 5.1. First-order metastases

The size *x* of a tumor of age *τ* is obtained by integrating the growth equation (Eq. (25)), from which we obtain

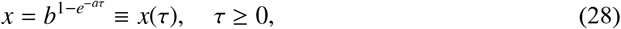

This expresses the size of the tumor as a function of its age. Solving for *τ* we obtain the age of the tumor as a function of size:

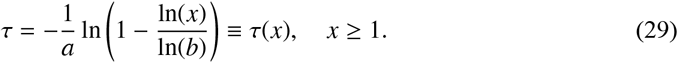

The two equations establish the equivalence between size and age. For the primary tumor we simply make the substitutions *x* = *x*_*p*_ and *τ* = *t* in the above equations. It follows that the emission rate of the primary tumor at time *t* is

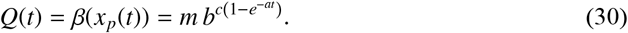

The first-order distribution of metastases, namely, metastases produced by the primary tumor, is given by Eq. (14). Combining Eq. (14) with the Gompertzian growth law (25) and the primary emission rate (30) yields

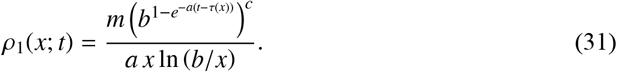

Thus we have the first-order solution of the metastatic model in analytic form.

### 5.2. Second-order metastases

The secondary emissions are calculated from Eq. (21) with *k* = 2:

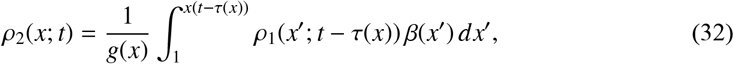

with *ρ*_1_(*x*; *t*) from (31) and *τ*(*x*) from (29). The integral must be evaluated numerically but the calculation is straightforward since everything on the righthand side is known. All integrals in the recursive formulation were evaluated numerically in MATLAB with adaptive quadrature method (function `integral`).

## 6. Results

The effect of the first three generations of metastatic tumors is shown in Figure 4 in years 1, 3, 4 and 5. Superimposed is the third-order approximant *ρ*^(3)^ = *ρ*_1_ + *ρ*_2_ + *ρ*_3_. The dynamic interplay between generations is not as simple as one might imagine. The dominant contribution to the distribution in year 1 comes from the first generation. The distributions of generations 2 and 3 have similar shapes but they are several orders of magnitude smaller. In this case the overall distribution is well approximated by the first-order approximant. In year 2 (not shown) all distributions increase and the curves in log–log scale move closer together over the size range where they overlap.

**Figure 4.**
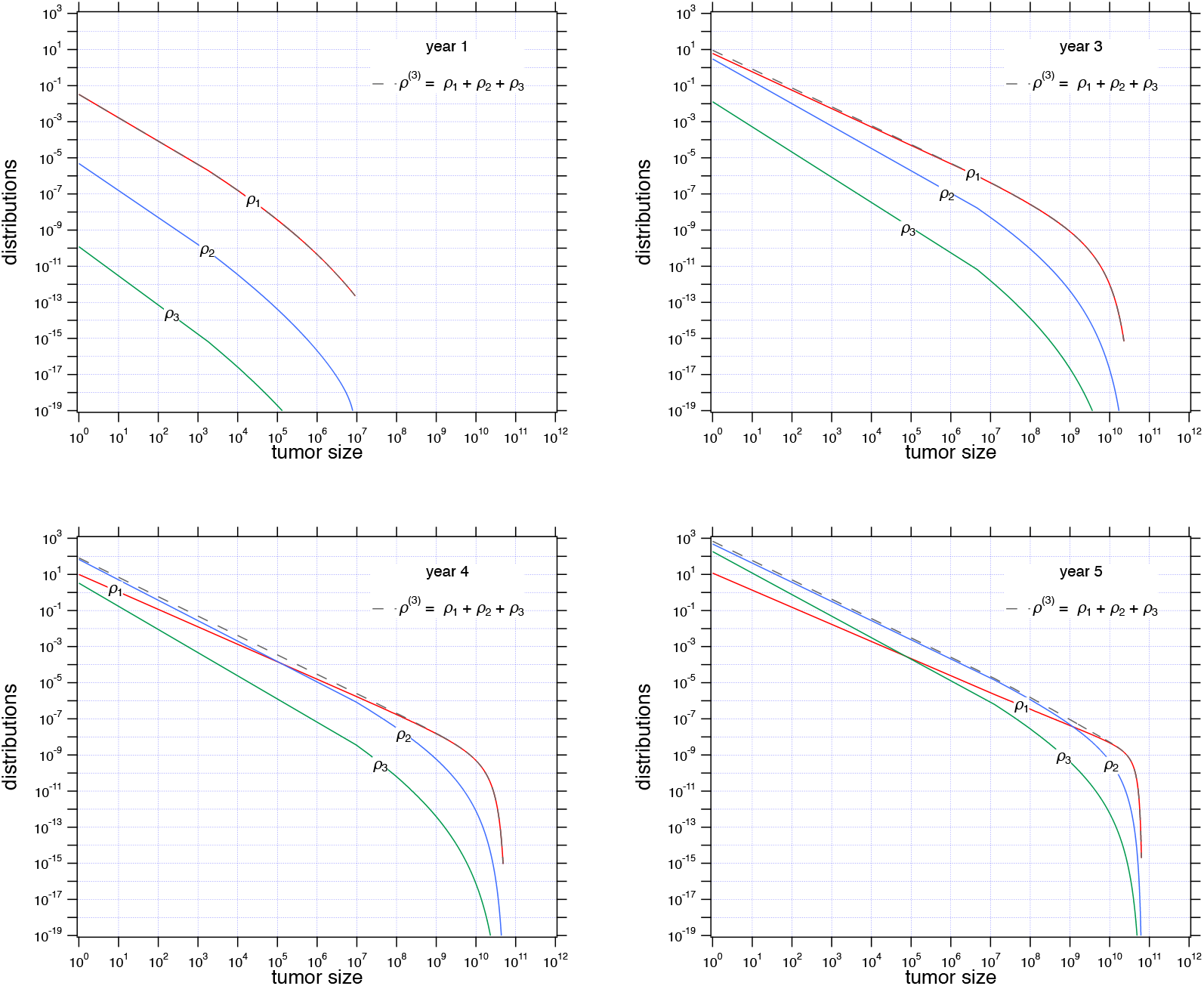
Distribution of tumors in generations 1, 2, and 3 at years 1, 3, 4, and 5. The plots show how different generations contribute over time: early on, the first generation dominates, but later, newer generations become increasingly significant, especially among smaller tumors.

By year 3, the distribution of the second generation begins to make a visible contribution in the lower size range (small *x*), while its impact at larger sizes remains limited. Generation 3 is more than two orders of magnitude smaller and makes no significant contribution. In year 4 the second generation overtakes the first at small sizes but not at the large ones. The third generation begins to make a contribution in the small range. As a result, the overall distribution at small sizes approximately follows the second generation while the tail follows the first generation. By year 5 it is the turn of the third generation to overtake the first generation at small scales. The general trend then is that younger generations initially grow slowly but eventually overtake older generations in the small size region. This is because the proliferation of tumors results in a commensurate proliferation of emissions. On the other hand, for each fixed time *t* and in the large-size region the distributions are ordered inversely by generation:

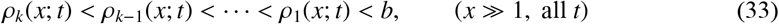

Eventually, the maximum tumor size—defined as the largest size reached among all tumors at time *t*—approaches the asymptotic value *x* = *b* dictated by the Gompertzian growth law. In other words, the right tail of the distribution *ρ*(*x*; *t*) moves toward *b* but never exceeds it.

The effect of metastatic generation is seen more clearly in the total number of metastatic tumors. We calculate the number of tumors in generation *k* by

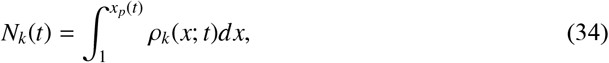

and the *k-*order approximation of the total number of tumors by

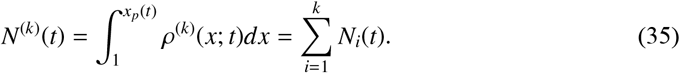

Figure 5a shows the number of metastatic tumors in the first two generations. The IKS model is characterized by an incubation period during which very few metastases are produced, due to the low emission rate from small tumors, followed by explosive growth once the size of the emitting tumors grows sufficiently. This behavior characterizes the emissions in all generations. Significantly, each subsequent generation has a longer incubation period than its parent generation, but also a more explosive growth. Until year 3 the primary emissions account for all metastatic cancers present; at about 3.75 years the secondary emissions match the primary emissions and quickly overtake them. In fact, the secondary emissions alone are a fairly good approximation of the total emissions because they overtake the effect of the primary emissions. By year 4 the third generation kicks in and can no longer be ignored. In Fig.5b we see the evolution of successive approximants. Each lower-order approximant begins at the true number of tumors but eventually underestimates that number as its offspring generation becomes large enough to produce significant number of metastases of its own.

**Figure 5.**
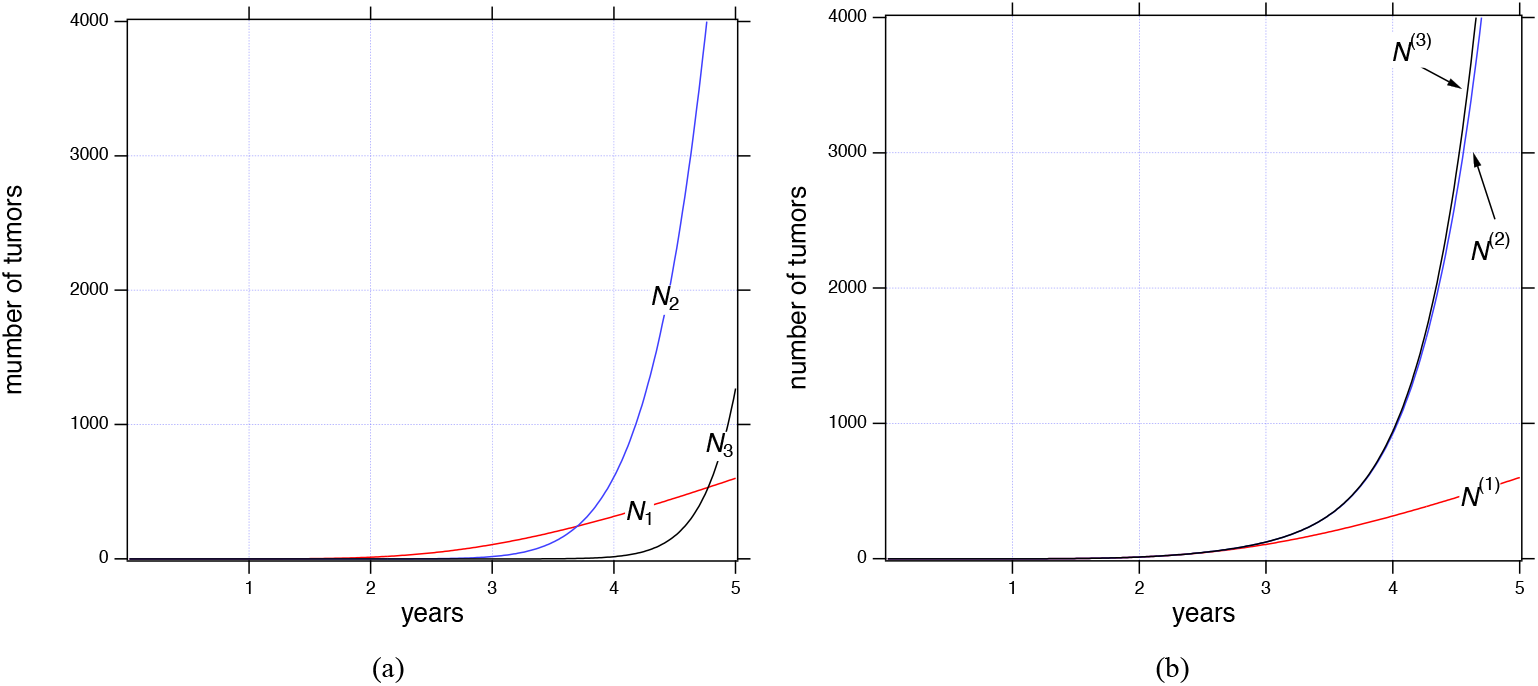
**Evolution of metastatic tumors in generations *N*_1_, *N*_2_ and *N*_3_** (a) and their cumulative totals *N*^(1)^, *N*^(2)^, and *N*^(3)^ (b) over a 5year period. Initially, each newer generation (e.g., *N*_2_ and *N*_3_) contributes fewer tumors, but as time progresses, they rapidly increase and eventually surpass the earlier generations, significantly raising the total metastatic burden. Note that *N*_*k*_(*t*) counts all metastatic foci, including microscopic deposits well below standard imaging thresholds; the number of radiologically detectable macrometastases is much smaller.

To address the quantitative interpretation of the model predictions, we now distinguish be tween the total number of metastatic foci (including microscopic seeds) and the number of macrometas-tases that would be biologically or clinically detectable. For a given threshold *x*_*t*_ (expressed in cells), we define the number of macrometastases as

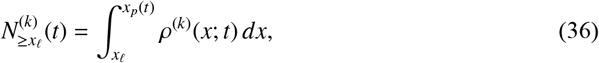

and the number of micrometastases (below the threshold) as the complement 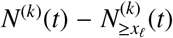. The threshold *x*_*𝓁*_ can be interpreted as the lower limit of clinical detectability, typically taken in the range 10^7^–10^9^ cells (corresponding to tumor diameters of about 3 mm 1 cm).

These patterns are illustrated in Figure 7, which shows the size-resolved distributions of metastases at two representative times (*T* = 3 and *T* = 3.8 years), separated by generation and stratified by the imaging threshold *x*_*t*_ = 10^8^ cells. For the baseline parameter set, the larger counts at later times (e.g. approximately *N*^(3)^(5 years) ≈ 4000 in Figure 5) therefore reflect predominantly sub-detectable micrometastases rather than thousands of macroscopic lesions. Quantitatively, the distributions in Figure 7 indicate that at *T* = 3 years the total number of metastatic foci is about *N*^(3)^(3) ≈ 125, of which only approximately 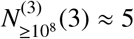 exceed the 10^8^cell threshold, leaving 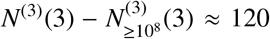 micrometastases below that threshold. By *T* = 3.8 years, the total burden has increased to approximately *N*^(3)^(3.8) ≈ 610, with 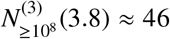 macrometas-tases above 10^8^ cells and 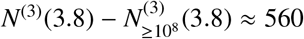 micrometastases. Thus, while the overall number of lesions grows by almost a factor of five between *T* = 3 and *T* = 3.8 years, the macroscopic component 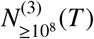 increases much more modestly, and higher generations contribute predominantly to the microscopic, subclinical pool.

While the number of tumors grows explosively with each generation, most of these tumors are young and therefore small. A more complete picture is obtained by looking at the mean size of the tumor. We calculate the *k*-order approximation of the mean tumor as

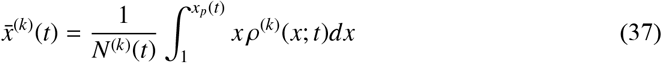

and plot in Fig. 6 the effect of the first three generations. The effect of each subsequent generation is to moderate the growth of the mean tumor size. Accordingly, a truncated approximant will eventually overpredict the size of the mean tumor but underpredict their total number.

**Figure 6.**
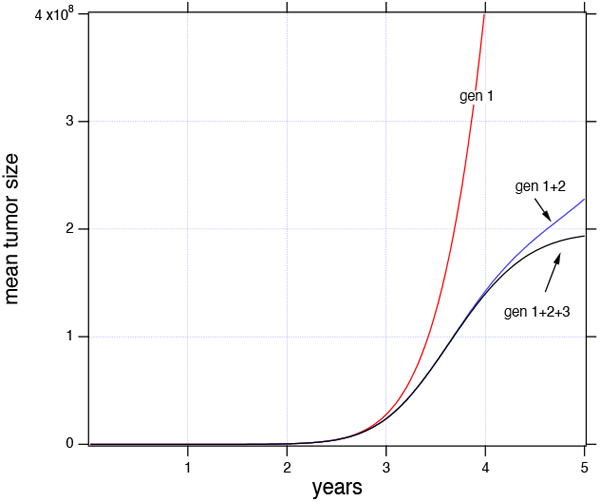
Mean tumor size over time, considering the cumulative effect of generations 1, 2 and 3. The plot highlights how adding newer generations (gen 1, gen 1 + 2 and gen 1 + 2 + 3) influences the average tumor size, demonstrating that accounting for younger generations reduces the overall mean tumor size.

As a final test we compare our results to the numerical solution of the IKS model by the method of characteristics (see Appendix C for numerical details). In logarithmic coordinates (Fig. 8a) the two solutions are in very good agreement over a five-year period.

**Figure 7.**
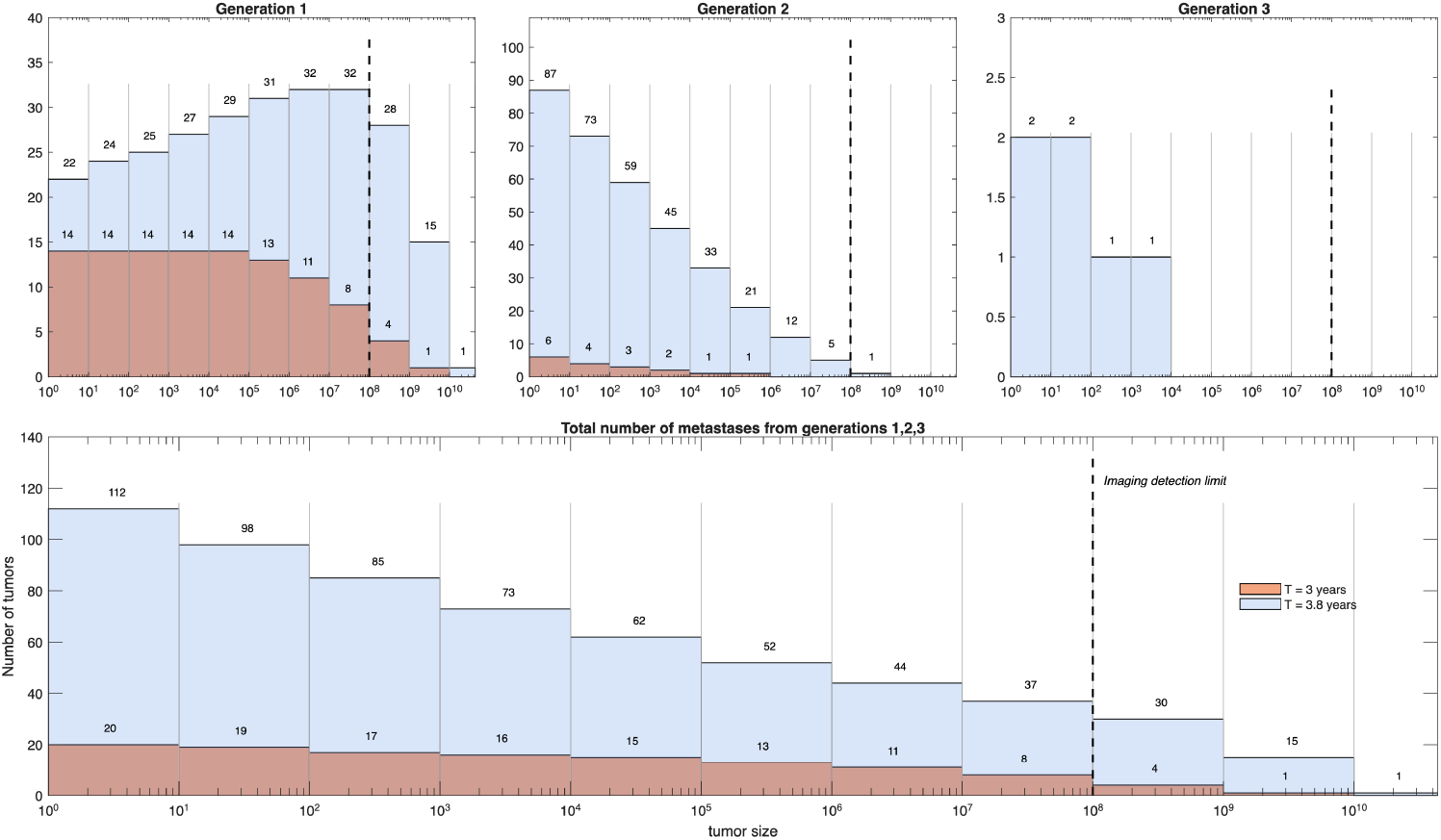
Size-resolved distribution of metastatic tumors at *T* = 3 years and *T* = 3.8 years, separated by metastatic generation (top panels) and shown cumulatively (bottom panel). Top: Histograms of tumor sizes for generations 1, 2, and 3. The dashed vertical line marks the imaging detection threshold (*x*_*t*_ = 10^8^ cells). Bottom: Cumulative number of tumors from generations 1–3 at *T* = 3 years (orange) and *T* = 3.8 years (blue). Bars to the right of the dashed line correspond to macrometastases (*x* ≥ 10^8^), while bars to the left represent micrometastases. Counts within each bin are indicated above the bars. The figure illustrates how successive generations contribute primarily to the microscopic tumor pool, with only a small fraction of tumors crossing the detection threshold, even as the total number of metastatic foci increases.

**Figure 8.**
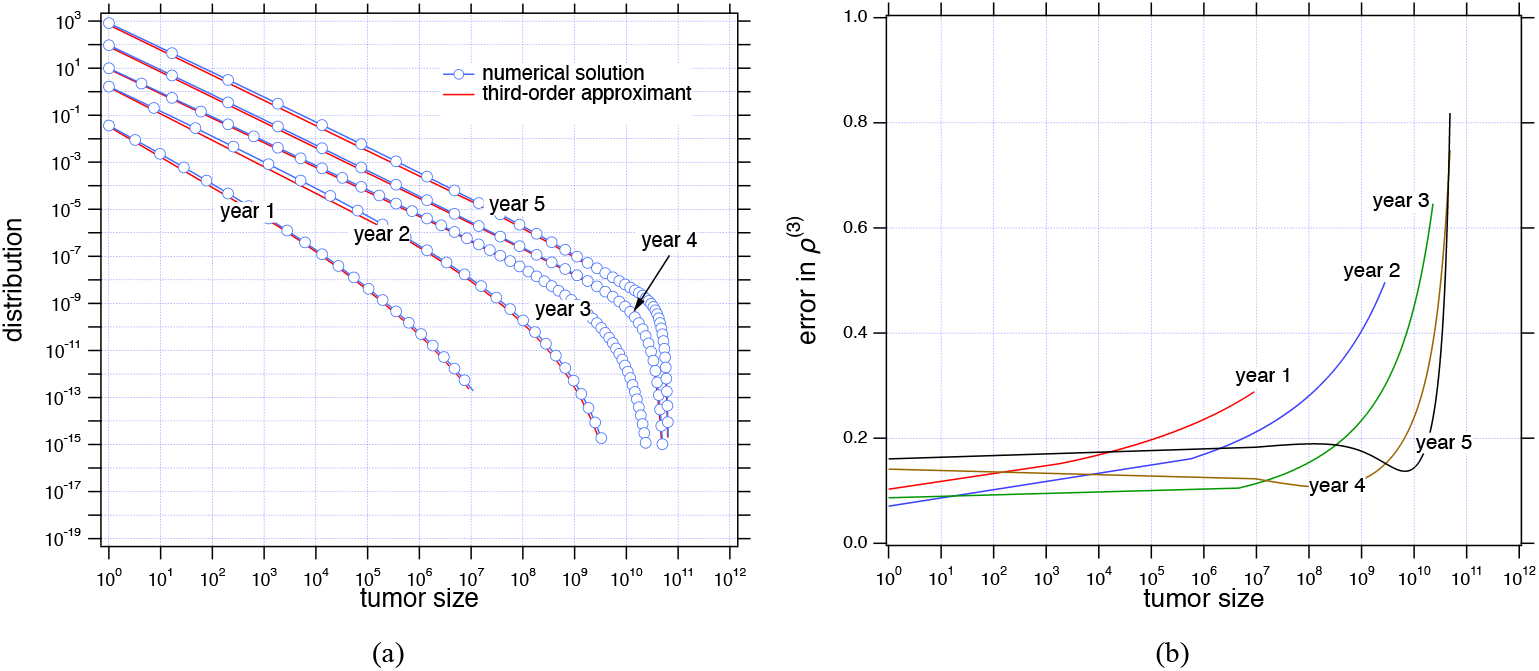
Comparison between the third-order approximant and the numerical solution of the IKS model over five years (a), along with their relative error |*ρ*^(3)^(*x*; *t*) − *ρ*_IKS_(*x*; *t*) | */ρ*_IKS_(*x*; *t*). (b). The plots demonstrate an agreement between the two approaches across tumor sizes, with minor deviations primarily observed at larger tumor sizes.

To make this comparison quantitative, we define the pointwise relative error in the size distribution as ❘ *ρ*^(3)^(*x*; *t*) − *ρ*_IKS_(*x*; *t*) ❘ */ρ*_IKS_(*x*; *t*) where *ρ*^(3)^(*x*; *t*) is the third-order recursive approximant and *ρ*_IKS_(*x*; *t*) is the numerical solution of the IKS transport equation obtained by the method of characteristics (Fig. 8b).

When we compute the relative error between the two methods the picture is somewhat less clear (Fig. 8b). The error is relatively constant over a wide range of tumor sizes up to about 10^6^. It grows at the tail of the distribution but this region makes insignificant contribution to the moments of interest. While the error in the region *x* 10^6^ remains constant, it ranges between 10% and 20%, higher than what we would expect between methods that converge to the same solution. We suspect that problem arises from the treatment of the boundary condition at *x* = 1. The balance on tumors of size *x* = 1 gives (see Appendix A)

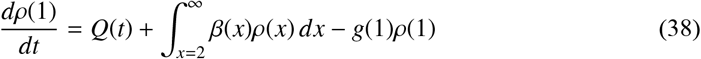

that reads: the number of tumors of size *x* = 1 increases by the rate of emissions from the primary tumor (given by *Q*(*t*)) plus the rate of emissions from all metastatic tumors (integral term), and decreases by the rate they grow out of size *x* = 1 (given by the term *g*(1)*ρ*(1)). To get from Eq. (38) the Eq. (3), which is the boundary condition of the IKS formulation, one must set *dρ*(1)*/dt* = 0. This can be justified if the condition

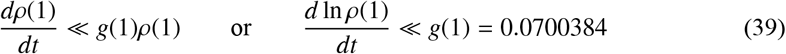

is true. This condition can be checked, since our methodology makes no use of the boundary condition.

The evolution of *ρ*^(3)^(1) is shown in Fig. 9a. It grows approximately as ~ *x*^4^, a very steep increase. Figure 9b shows the derivative *d* ln *ρ*^(3)^(1)*/dt* in comparison to *g*(1). Clearly, the condition in Eq. (39) is violated at all times. Accordingly, Eq. (3) is not the proper boundary condition for this model. This issue deserves further study.

**Figure 9.**
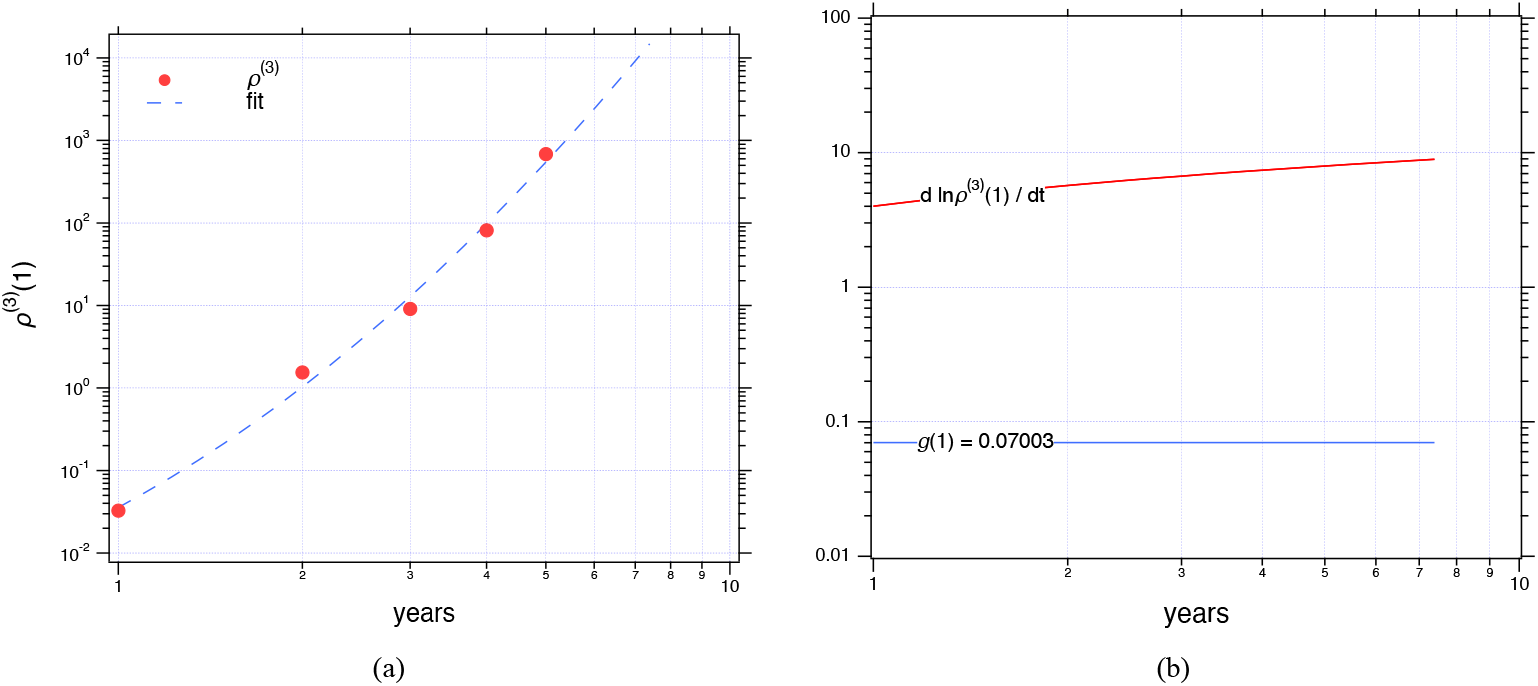
Analysis of the tumor density at the smallest tumor size (*x* = 1). Panel (a) shows how the density of tumors at this smallest detectable size increases dramatically over a 10year period, following a steep upward trend. Panel (b) provides a direct comparison between the rate at which this small-size tumor density changes over time, *d* ln *ρ*^(3)^(1)*/dt*, and the baseline tumor growth rate, *g*(1). This comparison reveals that the standard assumption used in modeling— specifically, that the density at the smallest size remains relatively stable—may not hold true, indicating the need for reconsideration of the boundary condition typically applied in metastatic tumor modeling.

## 7. Discussion

The aim the current work was to examine the consequences of exploiting the monotone correspondence between tumour size and age within the IKS framework[11]. By focusing on net growth and net dissemination, and lumping additional biological complexity—cell death, immune surveillance, colonisation failure and multistep extravasation and engraftment—into effective growth and emission rates *g*(*x*) and *β*(*x*), we obtained a deliberately minimal description of the metastatic cascade in which age and generation play the central roles. Mathematically, the key step is the replacement of the original size-structured PDE and its stiff nonlocal boundary condition by a recursive hierarchy of renewal-type integral equations indexed by metastatic generation. Along characteristics, tumour size and age are interchangeable state variables, and this observation yields closedform expressions for first-generation metastases and one-dimensional integral representations for higher generations.

From a numerical perspective, the renewal formulation avoids several fragilities of the standard PDE approach. Rather than enforcing a quasi-steady boundary flux at the minimal size *x*_0_ at every time step, the boundary flux emerges naturally from the history of growth and emissions encoded in the renewal kernels. The resulting recursion can be solved with standard quadrature schemes, without repeatedly solving a transport equation with a nonlocal boundary term. For the original IKS calibration to hepatocellular carcinoma [11], first-generation metastases seeded directly by the primary dominate both in number and in size during the early years of disease evolution; as time progresses, emissions from existing metastases generate second and third generations that overtake the first in lesion counts at small sizes, while older generations continue to populate the large-size tail. When an explicit detection threshold ( ≈ 10^7^ cells, below which metastatic foci are typically occult on standard CT, MRI, or PET imaging [36, 37]) is introduced, this generational separation naturally distinguishes a small number of clinically visible macrometastases from a much larger pool of microscopic foci.

A recurrent concern in interpreting the simulations is the absolute magnitude of total lesion counts at late times, but in our framework systemic burdens of order 10^3^–10^4^ foci by five years arise naturally in an aggressive, untreated scenario calibrated to a rapidly progressing clinical dataset. These figures should not be read as thousands of centimetre-scale tumours on imaging, but as the sum of microscopic and macroscopic lesions spanning several orders of magnitude in size. Once the distribution is truncated at a detection threshold comparable to CT/MRI resolution, the model predicts only a few to a few tens of macrometastases, consistent with advanced yet not exceptional presentations of aggressive disease. This interpretation is supported by multiple lines of evidence: miliary patterns in lung and liver consist of innumerable 1–5 mm nodules that are explicitly too many to count; Whole-organ clearing and light-sheet microscopy with DeepMACT-type pipelines reveal hundreds to thousands of micrometastases and single disseminated tumour cells per mouse, far below clinical imaging thresholds [38, 39]; rapid-autopsy programmes in metastatic breast cancer uncover far more lesions and a wider anatomical spread than premortem imaging suggests [40, 41]; and bonemarrow micrometastasis studies show that disseminated tumour cells are already present in a substantial fraction of early-stage patients and confer adverse prognosis [42, 43]. Within this empirical context, late-stage burdens of 10^3^–10^4^ predominantly microscopic foci are biologically plausible.

The current approach nevertheless still shares the same limitations as the original IKS model. In its simplest form, all lesions share the same growth and emission laws unless a generation dependent extension is imposed; organspecific microenvironments are ignored; and many mechanistic processes are absorbed into the effective net rates *g*(*x*) and *β*(*x*). Treatment, competition for resources and evolutionary changes in metastatic potential are not represented. Moreover, our analysis of the boundary flux at the minimal size suggests that the quasi-steady assumption underlying the standard IKS boundary condition is not satisfied by the recursive solution: the density of the smallest tumours increases too rapidly for *dρ*(*x*_0_; *t*)*/dt* to be neglected. This discrepancy merits further mathematical investigation and highlights the value of formulations in which the boundary condition is not imposed *a priori* but emerges from renewal considerations. Extensions in which growth and emission rates depend on metastatic generation, or microenvironment, in combination with spatial structure and Bayesian calibration against longitudinal imaging and molecular data, offer a natural route towards more realistic, patient-specific descriptions while retaining the conceptual clarity of the age–generation formalism.

## 8. Conclusions

In summary, we have recast the IKS model into an age and generation-based framework that replaces the original size-structured PDE and its nonlocal boundary condition with a family of solvable recursively equations across metastatic generations. Our reformulation remains fully consistent with Gompertz growth and power-law emission, reproduces characteristic-based numerical solutions of the IKS PDE over clinically relevant time scales, and is markedly simpler to analyse and numerically more stable. The generational perspective reveals a robust hierarchy in metastatic burden, with early disease dominated by lesions seeded directly from the primary and later stages dominated in number by younger, metastasis-derived generations, while older generations occupy the macroscopic tail. When an explicit detection threshold is imposed, the model naturally separates radiologically visible lesions from a much larger pool of microscopic foci and shows that, for the IKS hepatocellular carcinoma parameters, total burdens of order 10^3^–10^4^ lesions are compatible with realistic numbers of macrometastases and with existing pathological, autopsy and preclinical data. More broadly, the age–generation formalism offers a conceptually transparent and computationally efficient way to link primary-tumour growth, metastatic seeding across generations and the joint microscopic/macroscopic burden, thereby providing a quantitative explanation of why the metastases seen on imaging may represent only the visible tip of a substantially larger metastatic iceberg.

## CrediT authorship contribution statement

**Panagiotis Gavriliadis:** Writing – original draft, Methodology, Software. **Georgios Lolas:** Writing – original draft, Methodology, Funding acquisition. **Themis Matsoukas:** Writing – original draft, Methodology, Software, Conceptualization.

## Code Availability

The MATLAB programs implementing the recursive age–generation scheme and the numerical simulations are available from the corresponding author upon reasonable request.

## Acknowledgements

P.G. and G.L. would like to acknowledge funding from the European Innovation Council (EIC) under the European Union’s research and innovation program ALADDIN (grant agreement no. 101130574).

## Appendix A Population balance on metastatic tumors

Here we derive the continuous population balance equations (PBE) for metastatic cancer that form the basis of the mathematical treatments in the literature. We first obtain the discrete PBE and then pass to the continuous limit. The number *N*_*x*_(*t*) of tumors with *x* cells increases due to tumors with size *x* − 1 that grow to size *x*; decreases due to tumors of size *x* that grow to size *x* + 1; and decreases due to loss of one cell by metastasis. The balance equation reads

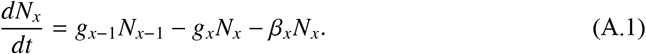

The equation for tumors with size *x* = 1 is different. The number of one-cell tumors increases by metastasis from the primary tumor at a rate *Q* and by metastasis from all metastatic tumors with rate

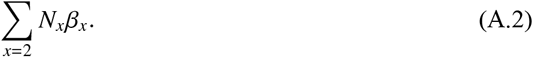

and decreases due to growth from size *x* = 1 to size *x* = 2. Therefore, the balance on the number of single-cell tumors is

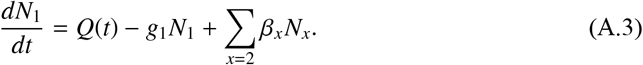

where *Q*(*t*) is the generation of rate of tumors from the primary tumor. Thus we have:

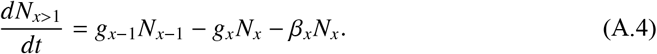

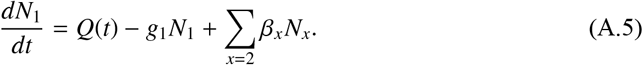

In the continuous limit (*x* » 1) we write *N*_*x*_ = *dN* = *ρ*(*x*)*dx* and the above equations become

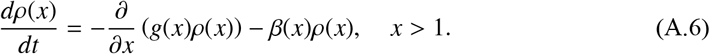

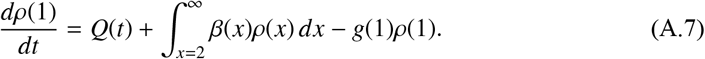

### Quasi Steady State

When the rate of metastasis is much smaller than the rate of growth (*b*(*x*) « *g*(*x*)) the term −*β*(*x*)*ρ*(*x*) may be dropped in Eq. (A.6) to obtain

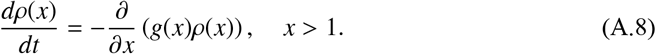

This is the form of the transport equation in the IKS mode.

Moreover, if one assumes that a quasi steady state is established at *x* = 1 such that the rate formation of single-cell tumors matches their rate of growth, then we may set *dρ*(1)*/dt* ≈ 0 and write the boundary condition in the form

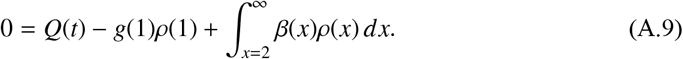

This is the boundary condition in the original IKS formulation as well as in all subsequent development. We note that our methodology does not require the quasi-steady state assumption and may be used to test directly the validity of Eq. (A.9), as explained in the text.

## Appendix B IKS-model: asymptotic solution

Iwata *et al* [11] obtained the analytic solution in the asymptotic limit *t* → ∞. It is given by:

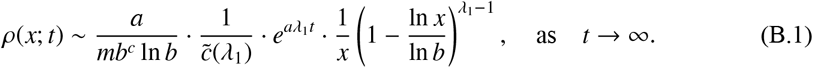

Where

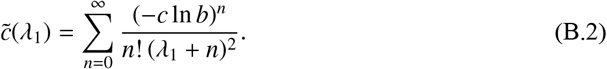

The parameter *λ*_1_ is the *unique positive solution* of the equation:

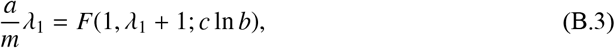

where *F* is the confluent hypergeometric function of Kummer, defined as:

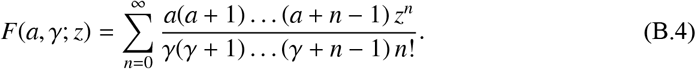

Equation (B.1) is of the form

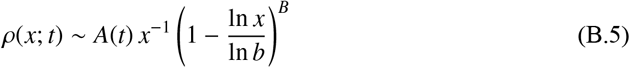

where *A* is a function of time but not of size and *B* is constant. Accordingly, for *x* « *b* the size distribution scales as *ρ*(*x*; *t*) ~ *x*^−1^. The distribution is truncated at *x* = *b*, which is the maximum possible tumor size in the Gompertz growth model.

## Appendix C IKS-model: method of characteristics

Here, we present the basic results of the integration of the IKS-model (Eqs. (2)–(3)) along the characteristic curves, introduced in [19] and implemented in the IKS-model by [15]. In this way, the integration of the PDEODE equations (Eq. (2) and Eq. (3)) is reduced to the integration of two ODE equations.

### The ODEs

We denote by 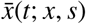 the characteristic curves, which are the solution of the initial value problem:

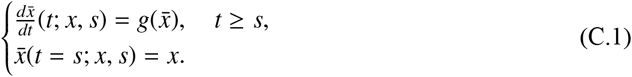

Thus, for the Gompertz function (see Eq.( (25)), the solution of the ODE (Eq. (C.1)) is explicitly given by:

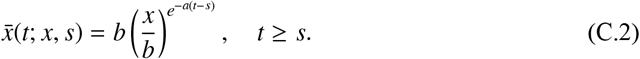

Along the characteristic curves 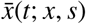, the transport equation Eq. (3) is reduced to the following ODE:

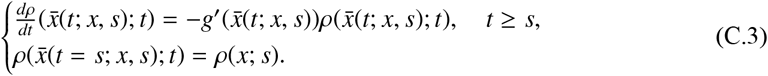

Thus, the solution of Eq. (C.3) along the characteristic curves Eq. (C.2) is given by:

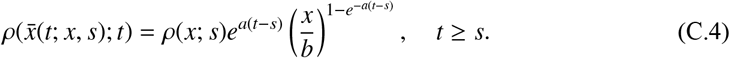

### Numerical scheme

Let the fixed time interval [0, *T*] and *k* a constant time step. We consider the uniform mesh in time interval with *k* = *T/N*:

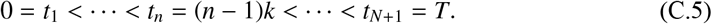

Then, we consider the associated spatial discretization in *x*, by means of the relation *x*_*n*_ = *x*_*p*_(*t*_*n*_) as follows:

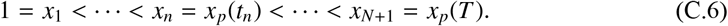

where *x*_*p*_(*t*_*n*_) is the primary tumor size at time *t*_*n*_. In this way, the uniform mesh in time interval [0, *T*] produces a nonuniform mesh in *x*-space [1, *x*_*p*_(*T*)].

Note that the resulting grid {(*x*_*i*_, *t*_*n*_), *i, n* = 1, .., *N* + 1} is constructed such that for every indices *i, n* the points (*x*_*i*_, *t*_*n*_) and (*x*_*i*+1_, *t*_*n*+1_) belong to the same characteristic curve (see [19]). Here, for completeness reasons we consider the solution algorithm of Barbolosi *et al* [15].

#### Algorithm 1

We denote with *ρ*^*n*^ (Eq. (C. 4)) an approximation of the solution of IKSmodel at the point (*x*_*i*_, *t*_*n*_).

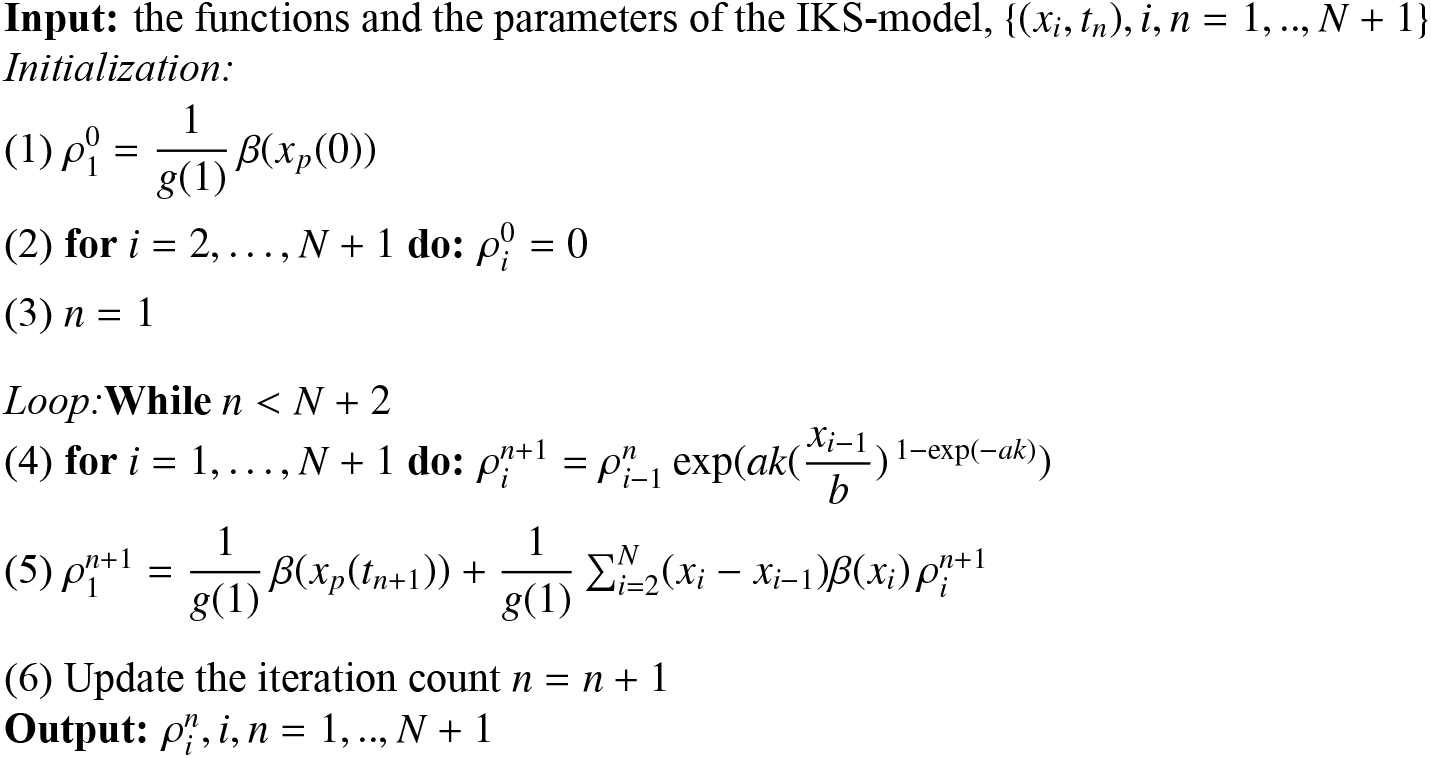

## Appendix D Nomenclature

**Table D.1.**
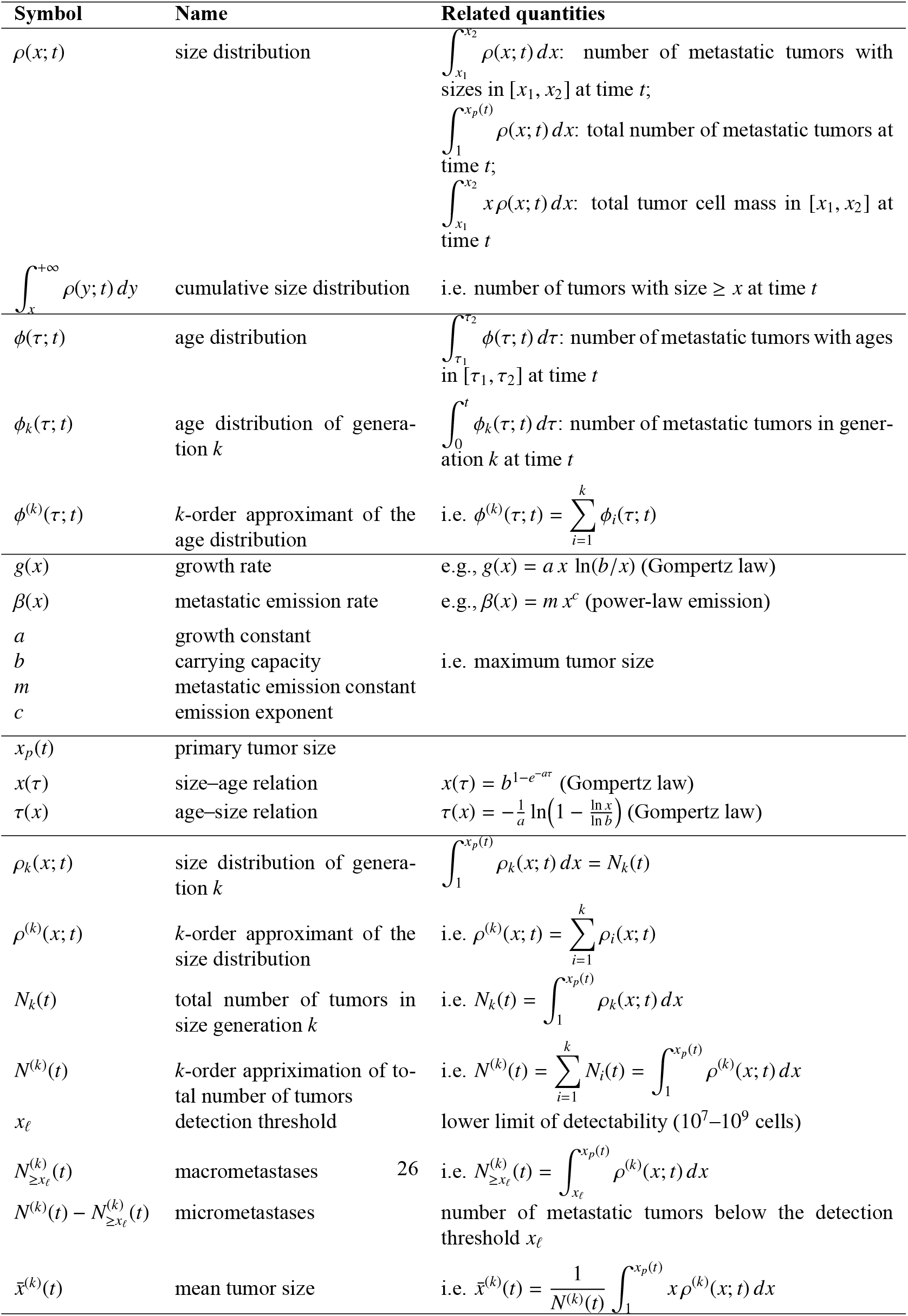
Nomenclature of main symbols and quantities.

## References

[1] W. Lambert, Y. Zhang, R. A. Weinberg, Cellintrinsic and microenvironmental determi nants of metastatic colonization, Nature Cell Biology 26 (2024) 687–697. 1

[2] A. Klein, Tumour cell dissemination and growth of metastasis, Nature Reviews Cancer 10 (2) (2010) 156–156. doi:10.1038/nrc2627-c2. URL http://dx.doi.org/10.1038/nrc2627-c2 1

[3] F. Chambers, A. C. Groom, I. C. MacDonald, Dissemination and growth of cancer cells in metastatic sites, Nature Reviews Cancer 2 (8) (2002) 563–572. doi:10.1038/nrc865. URL http://dx.doi.org/10.1038/nrc865 1

[4] S. Gerstberger, Q. Jiang, K. Ganesh, Metastasis, Cell 186 (8) (2023) 1564–1579. doi:10.1016/j.cell.2023.03.003. URL http://dx.doi.org/10.1016/j.cell.2023.03.003 1

[5] M. A. J. Chaplain, Avascular growth, angiogenesis and vascular growth in solid tumours: The mathematical modelling of the stages of tumour development, Mathematical and Computer Modelling 23 (6) (1996) 47 – 87. 1

[6] Ribba, N. Holford, P. Magni, I. Trocóniz, I. Gueorguieva, P. Girard, C. Sarr, M. Elish mereni, C. Kloft, L. Friberg, A review of mixed–effects models of tumor growth and effectsof anticancer drug treatment used in population analysis, CPT: Pharmacometrics amp; Sys tems Pharmacology 3 (5) (2014) 1–10. doi:10.1038/psp.2014.12. URL http://dx.doi.org/10.1038/psp.2014.12 1

[7] P. Altrock, L. Liu, M.F., The mathematics of cancer: integrating quantitative models, Nature Reviews Cancer 15 (2015) 730–745. 1

[8] Barbolosi, J. Ciccolini, B. Lacarelle, F. Barlesi, N. Andre, Computationaloncology mathematical modeling of drug regimens for precision medicine, Nature Reviews in Clinical Oncology 13 (2016) 242 – 254. 1

[9] A. M. Jarrett, E. A. B. F. Lima, D. M. A. Hormuth II, M. T. McKenna, X. Feng, D. A. Erkut, C. M. Resende, A. Brock, T. E. Yankeelov, Mathematical models of tumor cell proliferation: A review of the literature., Expert Rev Anticancer Ther. 18 (12) (2017) 1271–1286. 1

[10] C. Harkos, A. Hadjigeorgiou, C. Voutouri, F. Barlesi, A. Kumar, T. Stylianopoulos, R. Jain, Using mathematical modelling and ai to improve delivery and efficacy of therapies in cancer, Nature Reviews Cancer (2025). 1

[11] K. Iwata, K. Kawasaki, N. Shigesada, A dynamical model for the growth and size distribution of multiple metastatic tumors, Journal of Theoretical Biology 203 (2) (2000) 177–186. 1, 1.1, 1.1, 1.1, 5, 7, Appendix B

[12] A. G. McKendrick, Applications of mathematics to medical problems, Proc. Edinburgh Math. Soc. 44 (1926) 98–130. 1.1

[13] H. von Foerster, Some remarks on changing populations, in: J. F. Stohlman (Ed.), The Kinetics of Cellular Proliferation, Grune and Stratton, New York, 1959, pp. 382–407. 1.1

[14] M. Iannelli, Mathematical Theory of AgeStructured Population Dynamics, Applied Mathematics Monographs, Giardini editori e stampatori, Pisa, 1995. 1.1

[15] Barbolosi, A. Benabdallah, F. Hubert, F. Verga, Mathematical and numerical analysis for a model of growing metastatic tumors, Mathematical Biosciences 218 (1) (2009) 1–14. 1.1, Appendix C, Appendix C

[16] A. Devys, T. Goudon, P. Lafitte, A model describing the growth and the size distribution of multiple metastatic tumors, Discrete and Continuous Dynamical SystemsSeries B 12 (4) (2009) 731–767. 1.1, 1.3

[17] Stein, D. DeWoskin, M. Higley, K. Lemoi, B. Owens, A. Rahman, H. Rotstein, D. Rum schitzki, S. Swaminathan, M. Tanzy, O. Varfolomiyev, T. Witelski, V. Zubkov, Dynamic models of metastatic tumor growth, in: Final Report of the 27th Annual Workshop on Mathematical Problems in Industry, New Jersey Institute of Technology, 2011. 1.1

[18] J. Liu, X. S. Wang, Numerical optimal control of a sizestructured pde model for metastatic cancer treatment, Mathematical Biosciences 314 (2019) 28–42. 1.1, 1.2

[19] O. Angulo, J. C. LópezMarcos, Numerical schemes for sizestructured population equa tions, Mathematical Biosciences 157 (1-2) (1999) 169–188. 1.1, Appendix C, Appendix C

[20] N. Hartung, Efficient resolution of metastatic tumor growth models by reformulation into integral equations, Discrete and Continuous Dynamical Systems B 20 (2) (2015) 445–467. 1.1

[21] M. Bulai, M. C. De Bonis, C. Laurita, V. Sagaria, Modeling metastatic tumor evolution, numerical resolution and growth prediction, Mathematics and Computers in Simulation 203 (2023) 721–740. 1.1

[22] M. Bulai, M. C. De Bonis, C. Laurita, Numerical solution of metastatic tumor growth models with treatment, Applied Mathematics and Computation 484 (2025) 128988. 1.1

[23] P. Hahnfeldt, D. Panigrahy, J. Folkman, L. Hlatky, Tumor development under angiogenic signaling: A dynamical theory of tumor growth, treatment response, and postvascular dormancy, Cancer Research 59 (19) (1999) 4770–4775. 1.2

[24] S. Benzekry, Mathematical analysis of a twodimensional population model of metastatic growth including angiogenesis, Journal Evolution Equations 11 (2011) 187–213. 1.2

[25] S. Benzekry, A. Tracz, M. Mastri, R. Corbelli, D. Barbolosi, J. M. Ebos, Modeling spontaneous metastasis following surgery: an in vivoin silico approach, Cancer Research 76 (3) (2016) 535–547. 1.2

[26] Balbolosi, Modélisation du risque d’évolution métastatique chez les patients supposés avoir une maladie localisée, Oncologie 13 (2011) 528–533. 1.2

[27] M. Bilous, C. Serdjebi, A. Boyer, P. Tomasini, C. Pouypoudat, D. Barbolosi, F. Barlesi, F. Chomy, S. Benzekry, Quantitative mathematical modeling of clinical brain metastasis dynamics in nonsmall cell lung cancer, Scientific Reports 9 (1) (2019) 13018. 1.2

[28] Struckmeier, A mathematical investigation of a dynamical model for the growth and size distribution of multiple metastatic tumours, Niedersächsische Staats und Universitätsbib liothek, Göttingen OCLC:834703245 (2003) 1–13. 1.2

[29] Maiti, Monte Carlo simulationbased approach to model the size distribution of metastatic tumors, Physical Review E—Statistical, Nonlinear, and Soft Matter Physics 85 (1) (2012) 012901. 1.2

[30] N. Hartung, S. Mollard, D. Barbolosi, A. Benabdallah, G. Chapuisat, G. Henry, S. Giacometti, A. Iliadis, J. Ciccolini, C. Faivre, F. Hubert, Mathematical modeling of tumor growth and metastatic spreading: validation in tumorbearing mice, Cancer Research 74 (22) (2014) 6397–6407. 1.2

[31] P. Schlicke, C. Kuttler, C. Schumann, How mathematical modeling could contribute to the quantification of metastatic tumor burden under therapy: insights in immunotherapeutic treatment of nonsmall cell lung cancer, Theoretical Biology and Medical Modelling 18 (2021) 11. 1.2

[32] P. Schlicke, Mathematical modeling and lung cancer: quantifiable prognostic models improve clinical routine care, Ph.D. thesis, Technische Universität München (2022). 1.2

[33] A. A. Lesi, S. Heilmann, R. M. White, D. S. Rumschitzki, A new mathematical model for tumor growth, reduction and metastasis, validation with zebrafish melanoma and potential implications for dormancy and recurrence, BioRxiv (2019) 676791. 1.2

[34] M. Piraud, M. Wennmann, L. Kintzelé, J. Hillengass, U. Keller, G. Langs, M.A. We ber, B. H. Menze, Towards quantitative imaging biomarkers of tumor dissemination: A multiscale parametric modeling of multiple myeloma, Medical Image Analysis 57 (2019) 214–225. doi:10.1016/j.media.2019.07.001. URL http://dx.doi.org/10.1016/j.media.2019.07.001 1.2

[35] S. Benzekry, M. Mastri, C. Nicolò, J. M. Ebos, Machinelearning and mechanistic model ing of metastatic breast cancer after neoadjuvant treatment, PLOS Computational Biology 20 (5) (2024) e1012088. 1.2

[36] R. Y. Tsien, Imagining imaging’s future, Nature Reviews Molecular Cell Biology 4 (Suppl) (2003) SS16–SS21. 7

[37] P. S. Adusumilli, B. M. Stiles, M. Chan, D. P. Eisenberg, Z. Yu, S. F. Stanziale, R. Huq, R. J. Wong, V. W. Rusch, Y. Fong, Real-time diagnostic imaging of tumors and metastases by use of a replication-competent herpes vector to facilitate minimally invasive oncological surgery, The FASEB Journal 20 (6) (2006) 726–728. doi:10.1096/fj.05-5316fje. URL http://dx.doi.org/10.1096/fj.05-5316fje 7

[38] Pan, O. Schoppe, A. ParraDamas, R. Cai, M. I. Todorov, et al., Deep learning reveals cancer metastasis and therapeutic antibody targeting in the entire body, Cell 179 (7) (2019) 1661–1676.e19. doi:10.1016/j.cell.2019.11.013. 7

[39] TakahashiYamashiro, K. Miyazono, Tissue clearing method in visualization of cancer progression and metastasis, Upsala Journal of Medical Sciences 129 (S1) (2024) e10634. doi:10.48101/ujms.v129.10634. 7

[40] E. Avigdor, A. CiminoMathews, A. M. DeMarzo, J. L. Hicks, J. Shin, S. Sukumar, J. Fetting, P. Argani, B. H. Park, S. J. Wheelan, Mutational profiles of breast cancer metastases from a rapid autopsy series reveal multiple evolutionary trajectories, JCI Insight 2 (24) (2017) e96896. doi:10.1172/jci.insight.96896. 7

[41] T. Geukens, M. De Schepper, W. Van Den Bogaert, et al., Rapid autopsies to enhance metastatic research: the UPTIDER postmortem tissue donation program, npj Breast Cancer 10 (1) (2024) 31. doi:10.1038/s41523-024-00637-3. 7

[42] A. VincentSalomon, F.C. Bidard, J.Y. Pierga, Bone marrow micrometastasis in breast cancer: review of detection methods, prognostic impact and biological issues, Journal of Clinical Pathology 61 (5) (2008) 570–576. doi:10.1136/jcp.2007.046649. 7

[43] S. Braun, F. D. Vogl, B. Naume, et al., A pooled analysis of bone marrow micrometastasis in breast cancer, New England Journal of Medicine 353 (8) (2005) 793–802. doi:10.1056/NEJMoa050434. 7

